# Lipid specificity of the immune effector perforin

**DOI:** 10.1101/2020.04.22.054890

**Authors:** Adrian W. Hodel, Jesse A. Rudd-Schmidt, Joseph A. Trapani, Ilia Voskoboinik, Bart W. Hoogenboom

## Abstract

Perforin is a pore forming protein used by cytotoxic T lymphocytes to remove cancerous or virus-infected cells during immune response. During the response, the lymphocyte membrane becomes refractory to perforin function by accumulating densely ordered lipid rafts and externalizing negatively charged lipid species. The dense membrane packing lowers the capacity of perforin to bind, and negatively charged lipids scavenge any residual protein before pore formation. Using atomic force microscopy on model membrane systems, we here provide insight into the molecular basis of perforin lipid specificity.

## Introduction

Killer T cells or cytotoxic T lymphocytes (CTLs) kill virus-infected and cancerous cells to maintain immune homeostasis. During the immune response, CTLs form a synapse with their target cells in which they secrete the pore forming protein perforin and pro-apoptotic granzymes ^1,2^. Although both the CTL and target cell plasma membrane are locally exposed at the synapse to perforin, perforin forms oligomeric pores in the target cell membrane but not in the CTL ^3^. Through the pores, granzymes can enter and trigger apoptosis in the target cell [Figure 1]. By contrast, the CTLs remain impermeable to granzymes and thus remain viable, and can sequentially kill multiple target cells ^3,4^. Without such resistance, CTLs (as well as natural killer cells) would be as vulnerable to perforin as target cells. This would imply a (most costly) one-to-one ratio of killer cells to target cells and also prevent antigen experienced CTLs from differentiating into memory cells.

**Figure 1.**
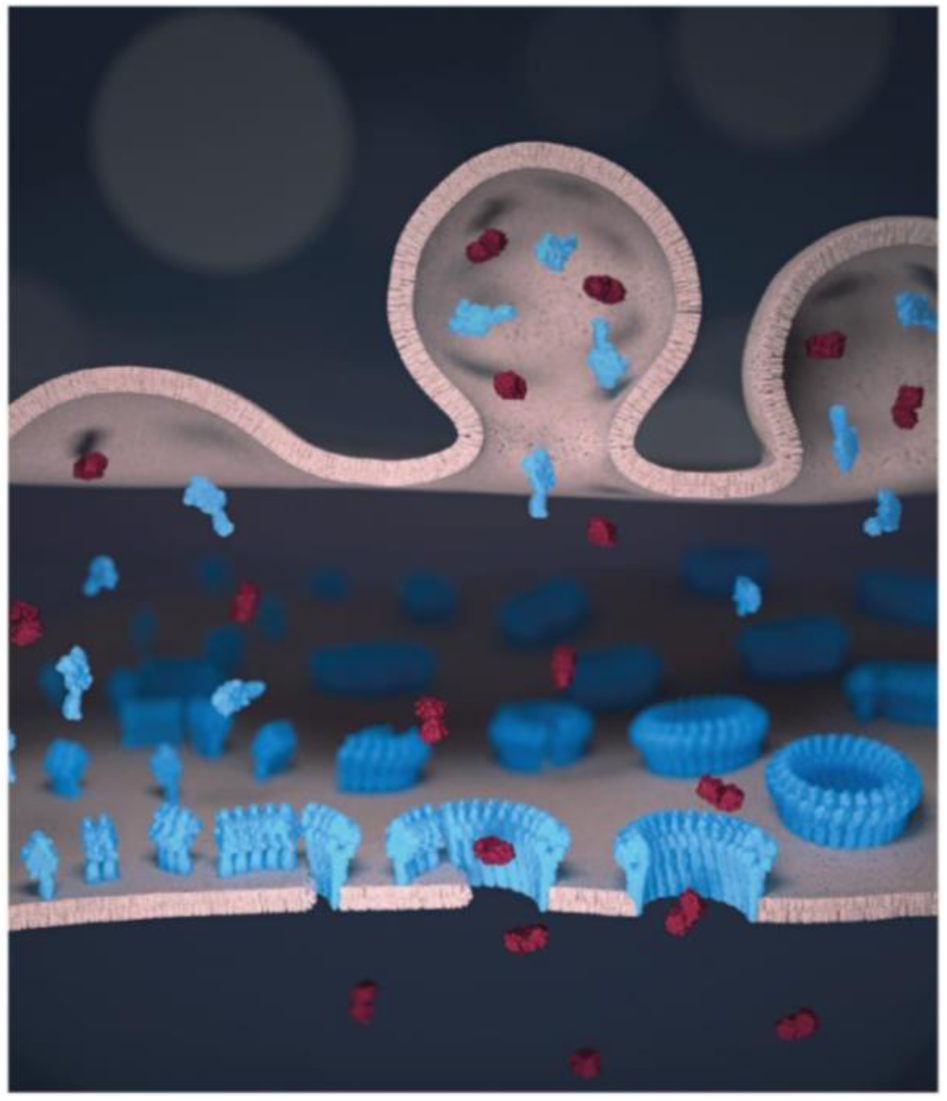
Schematic illustration of perforin pore formation and granzyme delivery in the synapse. Perforin (blue) and granzymes (red) are transported to the pre-synaptic membrane (top) by cytolytic granules and released into the synaptic cleft. The (monomeric) perforin subsequently binds to the target cell membrane (bottom). On the target membrane, from left to right, perforin first oligomerizes into short, non-lytic perforin prepores. Such prepores can convert to the pore state by inserting into the membrane, and subsequently recruit further prepores to sequentially grow the pore size. Once the pore size is sufficiently large, granzymes can diffuse into the target cells to trigger apoptosis.

Perforin membrane binding – the first step in pore formation - is calcium-dependent and is mediated by its C2 domain ^5–9^. It was initially thought that phosphocholine lipids were perforin receptors in the target membrane ^10^, but it was later shown – by a comparison of lipids well above and just below their gel transition temperature – that lipid order was a more important factor in determining membrane sensitivity to perforin^11^. The relatively tight plasma membrane packing of CTLs thus served as an explanation of the resistance of CTLs to perforin lysis. In the context of unidirectional killing in the immune synapse however, the hypothesis failed to explain the capability of CTLs to target and kill other CTLs, nor did it explain the absence of a clear correlation between the membrane packing of target cells and their susceptibility to perforin lysis ^12^. Following up on these early studies and on observations of perforin on model membranes by atomic force microscopy (AFM) ^13^, we have recently revealed a two-layered lipid-based mechanism that renders CTLs refractory to perforin pore formation ^14^: firstly, increased lipid order and packing in the CTL membrane reduces perforin binding to the membrane, and secondly, perforin is sequestered and irreversibly inactivated by binding to the negative charge of externalised phosphatidylserine (PS) at the CTL surface. Importantly, these membrane changes are enhanced in the area of the CTL plasma membrane that is associated with the immune synapse.

This lipid specificity can be regarded in the context of other pore forming proteins in general ^15^ and of the membrane attack complex-perforin/ cholesterol dependent cytolysin (MACPF/CDC) family of pore forming proteins, which perforin is part of ^16–19^. The bacterial CDCs use – as implied by their name – cholesterol as a receptor on the membrane and only form pores in membranes above a rather sharp threshold of cholesterol contents, typically above ∼25-35% ^20–23^. This cholesterol dependence defines the specificity of CDCs to eukaryotic target cells, as bacteria that generally do not contain cholesterol in their membranes ^24,25^. The mushroom derived MACPF pleurotolysin B utilizes the partner proteins ostreolysin A or pleurotolysin A to specifically bind and form pores in membranes containing sphingomyelin and cholesterol ^26–28^ or insect specific lipids ^29^. In vertebrates, the membrane attack complex (MAC) is an immune effector that kills pathogenic bacteria. The formation of the MAC is facilitated on membranes that contain negatively charged lipids and show increased membrane tension, mimicking the surface of Gram-negative bacteria ^30^. Other examples from the vertebrate immune system are the more recently discovered pore forming proteins of the gasdermin family, which share some structural elements with MACPF/CDCs and trigger cell death by perforating the membranes of infected cells from the inside out ^31^. Gasdermin pore formation is related to negatively charged lipids that are in the inner leaflets of eukaryotic plasma membranes and mitochondria.

Besides cell-based assays, lipid specificity of pore formation can be most conveniently studied on model lipid bilayers since these can be prepared from a wide selection of lipid components and therewith offer the ability to selectively alter biophysical properties. As a reference lipid, we used the dioleoyl derivative of phosphatidylcholine (PC), DOPC, which has a low liquid-gel transition, or melting, temperature (T_m_, ca. - 17 °C ^32^), and is therefore present in a liquid disordered (L_d_) state at physiological temperatures (37 °C). PC lipids in the L_d_ state (sometimes supplemented with cholesterol) are the most common components of model membranes used to visualize perforin assemblies ^7,13,14,33–35^, The addition of cholesterol to L_d_ membranes increases membrane order and, at sufficiently high concentrations, it can give rise to a liquid ordered (L_o_) state ^25^. Below their melting temperature T_m_, lipids exist in a solid ordered (S_o_) gel state. Different lipids have different T_m_, e.g., the dipalmitoyl derivative of PC, DPPC, has a T_m_ of ca. 41 °C ^36^ and is thus in the S_o_ state at physiological temperature. Over time, a membrane containing a mix of lipids can phase separate and display domains of different states of membrane order. Commonly used mixtures to mimic eukaryotic membranes use a low T_m_ lipid species like DOPC, cholesterol and high T_m_ sphingomyelin (SM), e.g., egg SM. In such mixtures, one readily observes phase-separation into PC-rich L_d_ domains and SM/cholesterol-rich L_o_ domains ^37,38^. Similarly, mixtures of DOPC and excess DPPC can lead to L_d_/S_o_ phase separation ^39^. The lipid phase state is an important factor to consider when mixing different types of lipids. Thus, to retain the L_d_ state of a reference DOPC bilayer, we can use the dioleoyl derivative of phospholipids, e.g. dioleoyl phosphatidylserine (DOPS, T_m_ ca. -11 °C ^40^), ethanolamine (DOPE, T_m_ ca. -8 °C ^41^), or glycerol (DOPG, T_m_ ca. -22 °C ^42^).

Such model membranes can be prepared as supported lipid bilayers on a flat substrate, e.g., mica or silica, facilitating their characterisation by in-liquid atomic force microscopy (AFM) experiments. AFM has become a popular tool to study mechanisms of pore forming proteins ^43,44^, in part because it allows a relatively straightforward distinction between prepore assemblies and membrane inserted pores. This can be achieved either by detecting a height change ^45,46^ or by the loss of mobility once the protein contacts the underlying substrate, respectively ^13,46^. Another important feature of AFM is its ability to distinguish between different lipid domains via Ångström-sized differences in membrane thickness ^47^, which allows to simultaneously detect lipid phase boundaries and protein pores.

Here we use AFM-based experiments to expand upon our recent work ^14^ on establishing and elucidating how perforin function depends on the physicochemical properties of target membranes. Noting that perforin binding – and thus pore formation – is reduced on tightly packed membranes, we study and compare the effects of several properties that modulate membrane packing. In contrast to its response to changes in membrane order/packing, the interaction of perforin with the negatively charged DOPS is fundamentally different. The initial binding of perforin appears to be unaffected, but pore formation is disrupted. On model membranes, this effect is proportional to the amount of DOPS they contain. We therefore investigate how perforin interacts with pure DOPS membranes in the pursuit of understanding how perforin is deactivated by this lipid. Lastly, we describe the interaction of perforin with DOPE, as PE lipids are another major constituent of the plasma membrane.

## Experimental

### Recombinant proteins

Wild-type perforin (WT-PRF)^48^, disulphide locked perforin (TMH1-PRF)^13^, GFP fusion disulphide locked perforin (TMH1-GFP-PRF)^14^, C2 domain mutant perforin (D429A-PRF)^5^ were expressed in baculovirus-infected Sf21 cells and purified from the supernatant as per the respective references provided. The CDC perfringolysin O (PFO) was kindly provided by Rana Lonnen and Peter Andrew (University of Leicester).

### Preparation of lipid vesicles and AFM samples

1,2-dioleoyl-sn-glycero-3-phosphocholine (DOPC), 1,2-dipalmitoyl-sn-glycero-3-phosphocholine (DPPC), 1,2-dioleoyl-sn-glycero-3-phosphoethanolamine (DOPE), 1,2-dioleoyl-sn-glycero-3-phospho-(1’-rac-glycerol) (DOPG), 1,2-dioleoyl-sn-glycero-3-phospho-L-serine (DOPS), egg sphingomyelin (egg SM) and cholesterol were purchased as powder from Avanti Polar Lipids (Alabaster, AL, USA). Where indicated, the lipids were mixed in the desired molar ratios (±5% confidence intervals). At a concentration of 0.5-1 mg/mL, small unilamellar vesicles with a nominal diameter 100 nm were prepared using the lipid extrusion method ^13,49^.

4-8 µL of the small unilamellar vesicles (containing 4 µg of lipid) were added to a freshly cleaved, ø10 mm mica disc (Agar Scientific) topped up with 80 µL of adsorption buffer containing 150 mM NaCl, 25 mM Mg^2+^, 5 mM Ca^2+^, 20 mM HEPES, pH 7.4. To form a pure DOPG bilayer, lower salt conditions were necessary ^50^ and the buffer was thus adjusted, instead containing no Mg^2+^ and 10 mM Ca^2+^. The lipids were incubated for 30 minutes above the T_m_ of constituent lipids to cover the mica substrate with an extended lipid bilayer. Excess vesicles were removed by washing the bilayer 6-12 times with 80 µL of the adsorption buffer.

Additional washed were applied to samples that contained DOPS, where we found that Mg^2+^ in the buffer interferes with perforin binding, that contained DOPG to remove excess Ca^2+^, or to control samples that require the removal of Mg^2+^: these were washed an additional 6 times with 80 µL buffer containing 150 mM NaCl, 5 mM Ca^2+^, 20 mM HEPES, pH 7.4.

Wild-type perforin (WT-PRF), disulphide-locked perforin (TMH1-PRF), C2 domain mutant perforin (D429A-PRF) and perfringolysin O (PFO) were diluted up to ca. ten-fold to a volume of 40 µL in 150 mM NaCl, 20 mM HEPES, pH 7.4, and injected onto the sample, to a reach concentrations of 150 nM or, where noted, ca. 400 nM above the model membrane. The protein was incubated for 2 (where noted) or 5 min at 37 °C. To unlock TMH1-PRF after its binding to the membrane, the engineered disulphide bond was reduced by addition of 2 mM DTT (Sigma-Aldrich) and 10 min incubation at 37 °C. We previously verified that, once TMH1-PRF was bound to target membranes, the effect of DTT on its native disulphide bonds did not change the pore forming functionality ^13^.

### AFM imaging and analysis

AFM images were either recorded by force-distance curve-based imaging (PeakForce Tapping) on a MultiMode 8 system (Bruker, Santa Barbara, CA, USA) or photothermal excitation (blueDrive) on a Cypher ES AFM (Oxford Instruments, Abingdon, UK). The imaging conditions with commercial MSNL cantilevers (Bruker) for PeakForce Tapping are outlined in Ref. ^14^. In brief, PeakForce Tapping was performed at 2 kHz and a maximum tip-sample separation of 5-20 nm. Images were recorded at 0.75 Hz scan speed and tip-sample interaction forces between 50 and 100 pN on an E-Scanner (Bruker, Santa Barbara, CA, USA) with temperature control. For blueDrive, we used BL-AC40TS probes (Olympus, Tokyo, Japan). The UV laser for photothermal excitation was focussed onto the cantilever base. The laser was tuned to the resonance frequency of the cantilever in liquid (ca. 25 kHz) and the amplitude adjusted to 1 V. Imaging was performed at an amplitude setpoint of ca. 750 mV and 1 Hz scan speed. All samples were imaged at 37 °C to retain thermotropic properties, or at room temperature.

Raw AFM images were background subtracted with reference to the lipid surface, masking perforin and applying second-order flattening. Height values indicated in the manuscript are given with ±1 nm confidence intervals, with the uncertainty due to scanner calibration and possible sample deformation caused by the probe-sample interaction. The same colour/ height scale was applied to all images, spanning 25 nm and 9 nm below and 16 nm above the membrane surface (set to 0 nm). The colour scale is only depicted once, in the first figure. Values for perforin coverage were either estimated by the area above a height threshold located 6-8 nm above the membrane surface and adjusted to counteract tip broadening effects; or, when sufficient images at a higher pixel resolution were available, by tracing pore shapes with 3dmod 4.9.4 (BL3DEMC & Regents of the University of Colorado ^51^). Perforin coverages obtained by both methods are normalized with respect to a 100% DOPC reference and given as values between 0 and 1. One-way analysis of variance (ANOVA) with Dunnett’s post-hoc analysis was performed in R-3.6.3 using the multcomp package ^52^.

### Perforin binding to lipid strips

Membrane Lipid Strips (Echelon Biosciences, Salt Lake City, UT, USA) were incubated in 4 mL of blocking buffer containing 3% w/w BSA (Roche Diagnostics GmbH, Mannheim, Germany), 150 mM NaCl, 20 mM HEPES, pH 7.4 for 1 hr at room temperature. 2 µg/mL TMH1-GFP-PRF was added to a lipid strip in 4 mL of blocking buffer supplemented with 2 mM CaCl_2_, pH 7.4. The use of the GFP fusion construct allowed readout of the lipid strips without the need for antibody labelling. To assess calcium-independent (non-specific) perforin binding, 2 µg/mL TMH1-GFP-PRF in 4mL of blocking buffer was added to a lipid strip. After 1 hr of incubation at room temperature, the lipid strips were washed three times with 4 mL of blocking buffer (with or without added 2 mM CaCl_2_ to match the initial incubation). GFP fluorescence (of wet lipid strips) as a measure of perforin binding was recorded on an iBright 1500 western blot imaging system (Thermo Fisher Scientific, Waltham, MA, USA). The strips were stored in the blocking buffer for ca. 96 hours at 4 °C and imaged again.

## Results and discussion

### Effect of lipid order on perforin binding and pore formation

To visualize binding of perforin to different phase domains of different levels of lipid order, we used a disulphide-locked mutant, TMH1-PRF, that can bind to and assemble on, but not insert into the target membrane ^13^. Its pore-forming functionality was fully restored after reducing the disulphide bond with a reducing agent dithiothreitol (DTT). This mutant has the advantage over the wild-type protein in that membrane binding and pore insertion can be uncoupled and studied as two separate events ^13,14^.

Using TMH1-PRF, we first validated the earlier observation ^11^ that perforin does not bind to lipids that are below their gel-transition temperature, *i.e*., in the gel or S_o_ phase. This could be made most articulate by visualizing the binding of perforin on phase separated bilayers that contained both L_d_ and S_o_ domains. To this end, we mixed high T_m_ DPPC and low T_m_ DOPC and verified phase separation by AFM, with the S_o_ domains appearing ca. 1 nm higher than the L_d_ domains, see [Figure 2]. Upon exposure to TMH1-PRF, L_d_ domains showed extensive protein coverage by the appearance of diffuse plateaus with a height close to 10 nm above the membrane. Previously, such plateaus were associated with the membrane-bound but not inserted and, hence, highly mobile perforin prepores ^13^. After addition of DTT, the TMH1-PRF transited from the prepore to the membrane-inserted pore state, and the pores were only found on L_d_ domains.

**Figure 2.**
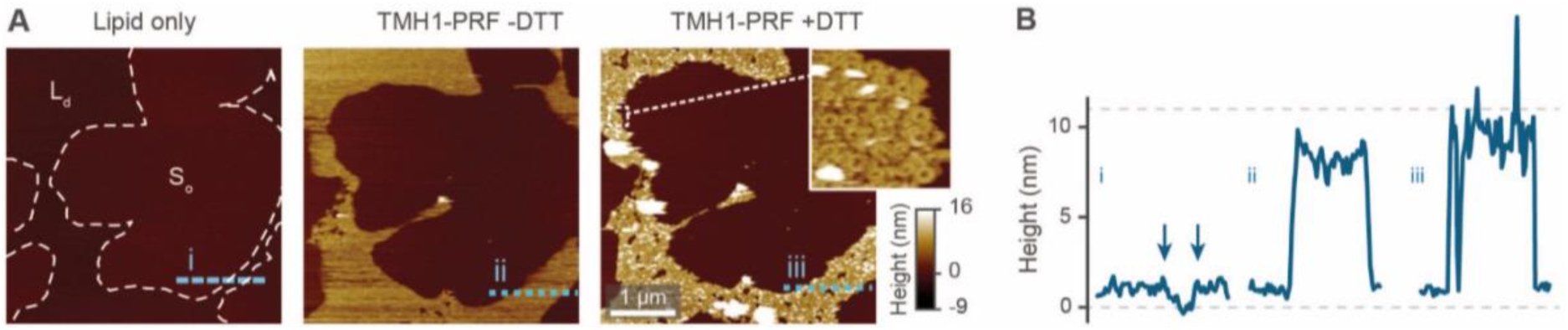
Prepore-locked TMH1-PRF preferentially binding to L_d_ domains in a phase separated L_d_/S_o_ membrane. (A) AFM images of a supported lipid bilayer composed by DOPC/DPPC mixed in a 1:7 molar ratio. The first panel (‘Lipid only’) shows the empty membrane, with the phase boundaries outlined by dashed white lines. Addition of TMH1-PRF (‘TMH1-PRF -DTT’) leads to the formation of a diffuse plateau limited to L_d_ domains. Similar plateaus were earlier interpreted as mobile prepore assemblies ^13^. After addition of DTT (‘TMH1-PRF +DTT’), the mobile assemblies insert into the membrane, and a dense layer of arc- and ring-shaped pores is formed (see inset), still confined to L_d_ domains. Size of the inset, 150 nm. (B) Height profiles extracted along the dashed coloured lines in A. The profiles depict the 0.5-1 nm height change at phase boundaries (i, the boundaries are highlighted by arrows), and the ca. 7-11 nm tall prepore and pore layers (ii and iii). Dashed grey lines indicate the height of the Ld membrane (0 nm) and the height of a perforin monomer (11 nm ^7^). Note that perforin features can appear compressed due to tip-sample interaction forces. The data was recorded at 37 °C.

Besides bilayer-based experimental substrates, membrane strips blotted with different types of lipids have been used to characterize lipid specificity of several pore forming proteins (e.g. ^29,53–55^). We find that commercially available strips fail to detect perforin binding to, for example, PC [Figure 3], in agreement with previous reports ^54 §^; this is in apparent contradiction with the scientific literature spanning from the 1980s ^10^ to today ^14^. This contradiction can be simply explained by noting that the blotted (phospho-) lipids have a high T_m_ and were not in an L_d_ state at the physiological/ experimental temperature, and are thus likely to cause erroneous readings when lipid order is a factor of importance for protein binding (such as for perforin). Moreover, lipids that bound perforin on such lipid strips ([Figure 3B,C, ‘+Ca^2+^’ vs. ‘-Ca^2+^’]) did so independently of the calcium that is required for perforin binding to target membranes; such interactions may therefore be artefacts due to defects in the lipid covered membrane surfaces.

**Figure 3.**
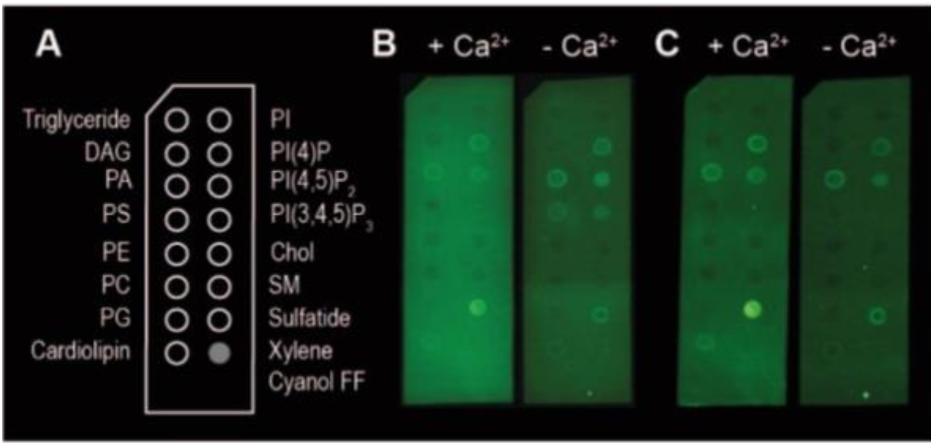
Binding of TMH1-GFP-PRF to lipid strips in the presence and absence of 2 mM Ca^2+^, visualized by fluorescence imaging ^§^. (A) Schematic layout of the lipid strips, with spots of lipids blotted where indicated (Xylene Cyanol FF is a non-lipid control). (B) Detection of TMH1-GFP-PRF binding to lipid strips in the presence and absence of Ca^2+^ (‘+ Ca^2+^’ and ‘-Ca^2+^’ respectively) immediately after washing. (C) The same lipid strips as in B after 4 days at 4 °C in blocking buffer.

As reported previously ^14^ and reiterated here for completion and for comparison with [Figure 2], we observe the same preference of perforin for L_d_ domains when adding TMH1-PRF to phase separated lipid membranes that contain both L_d_ and L_o_ domains ^14^, also in agreement with previous results using WT-PRF ^13,14^. As shown in [Figure 4], perforin again preferentially binds to and forms pores in L_d_ domains, albeit that some rare examples of perforin binding may be observed on L_o_ domains. This binding dependence on lipid order is also in agreement with the reduction of WT-PRF pore formation on highly ordered egg sphingomyelin membranes compared with L_d_ 18:1 sphingomyelin membranes ^14^.

**Figure 4.**
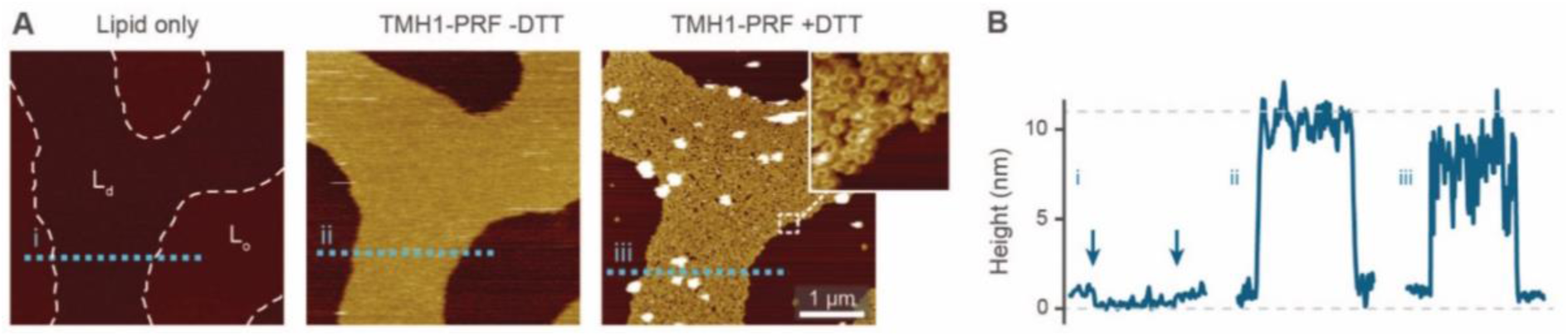
TMH1-PRF preferentially binding to L_d_ domains in a phase separated L_d_/L_o_ membrane, analogous to Figure 2. (A) AFM images of an approximately equimolar DOPC/egg SM/cholesterol supported lipid bilayer. ‘Lipid only’ shows the empty membrane with the phase boundaries between L_d_ and L_o_ domains highlighted by dashed white lines. TMH1-PRF exclusively binds L_d_ domains (‘TMH1-PRF -DTT’) and remains (mostly) confined there after addition of DTT (‘TMH1-PRF +DTT’) and the formation of transmembrane pores (see inset). Size of the inset, 150 nm. (B) Height profiles extracted along the dashed coloured lines in A. The profiles depict the 0.5-1 nm height change at phase boundaries (i, the boundaries are highlighted by arrows), and the ca. 7-11 nm tall prepore and pore layers (ii and iii). Dashed grey lines indicate the height of the L_d_ membrane (0 nm) and the height of a perforin monomer (11 nm ^7^). Note that perforin features can appear compressed due to tip-sample interaction forces. The data was recorded at 37 °C. Figure reproduced from Ref. ^14^, permission pending.

In the experiments reported above, lipid order was varied by using lipids of identical headgroups but different hydrophobic tails. In addition, membrane order can be dependent on divalent ions that intercalate with lipid headgroups, modulating intermolecular attractions ^56^. In most of our AFM work on model membranes, we had used up to 25 mM of Mg^2+^ in our buffers to stabilize supported lipid bilayers on the negatively charged mica substrate. This concentration is about one order of magnitude higher than blood levels ^57^. To test how the presence of Mg^2+^ affects perforin function, we designed experiments in which we washed samples to remove Mg^2+^ from the buffer before adding perforin (WT-PRF, at 37 °C as usual) onto model membranes either in the L_d_ (pure DOPC), L_o_ (DOPC/cholesterol or egg SM/cholesterol, both 47/53 molar ratio), or S_o_ (pure egg SM) state ^37^.

By comparison with previously published data acquired in the presence of Mg^2+^, we found the differences between pore formation at high and low levels of Mg^2+^ to be mostly insignificant, see Figure 5. However, for low levels of Mg^2+^, a significant but a small increase in pore formation was found on the L_o_ membranes consisting of DOPC/cholesterol and egg SM/cholesterol. We did not note any phase separation in any of these lipid substrates at either level of Mg^2+^, suggesting that perforin binding was uniformly affected, if at all, in all samples. In summary, a suggested increase in membrane order due to Mg^2+^ may be present and affect perforin pore formation in membranes of intermediate order, but this effect is small compared to the effects on lipid order due to high amounts of cholesterol or introduction of gel-phase lipids as reported above.

**Figure 5.**
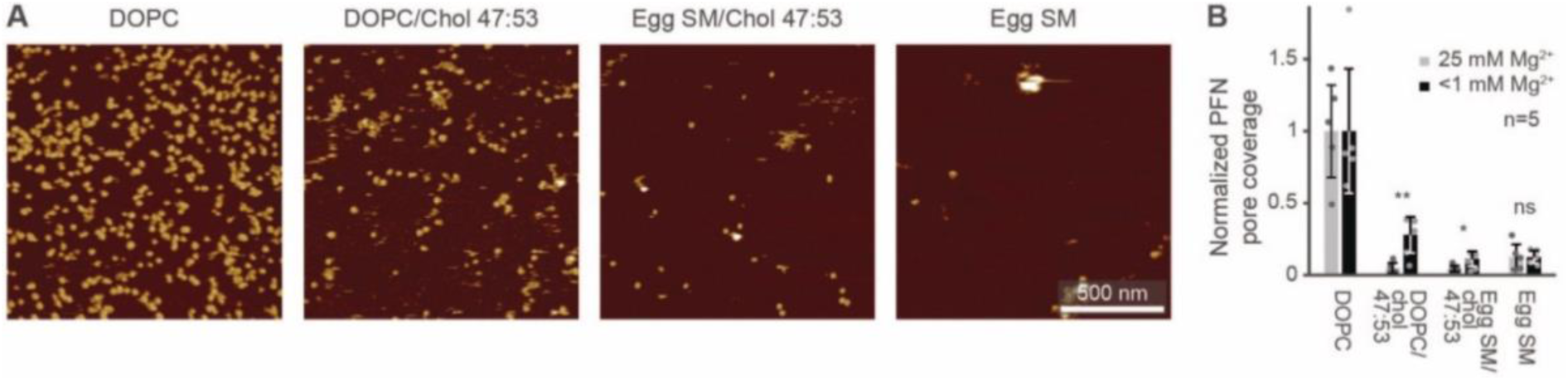
Perforin lipid specificity as a function of Mg^2+^ concentration. (A) Representative AFM images of WT-PRF pores on magnesium-depleted membranes (see Methods) of different lipid compositions, as indicated. The data was recorded at 37 °C. (B) Average perforin pore formation on different lipid mixtures, at Mg^2+^ concentration of 25 mM and <1 mM, normalized to the number of pores on DOPC-only membranes. Here, perforin was incubated for 2 min instead of 5 min (see Methods) to match experimental conditions of the two datasets. Error bars represent standard deviations. The statistical significance was assessed using ANOVA with Dunnett’s post-hoc analysis, where ‘ns’ is not significant, * p < 0.05, ** p < 0.01. Data for 25 mM Mg^2+^ are reproduced from Ref. ^14^.

### Effect of lipid charge on perforin pore formation

By the here described variations in perforin binding with lipid order, we can explain the reduced binding of perforin to killer lymphocytes. However, when incubated with higher concentrations of recombinant perforin, lymphocytes were found to still resist perforin pore formation in spite of binding amounts of perforin that were lytic to target cells, which we attributed to the presence of PS in the outer leaflet of the lymphocyte membranes ^14^.

Perforin can bind to PS-rich membranes, but pore formation is decreased: instead of pores, perforin aggregates into dysfunctional plaques ^14^. PS lipids have a net negative charge at physiological pH, and we previously hypothesized that this negative charge is the underlying cause of perforin dysfunction. We therefore tested the effect of the negatively charged DOPG and cholesterol sulphate on perforin pore formation. As predicted, the decrease in perforin pore formation was proportional to the levels of negatively charged lipids in the membranes and, possibly, further affected by an ordering effect induced by cholesterol sulphate (Figure 6 A,B).

**Figure 6.**
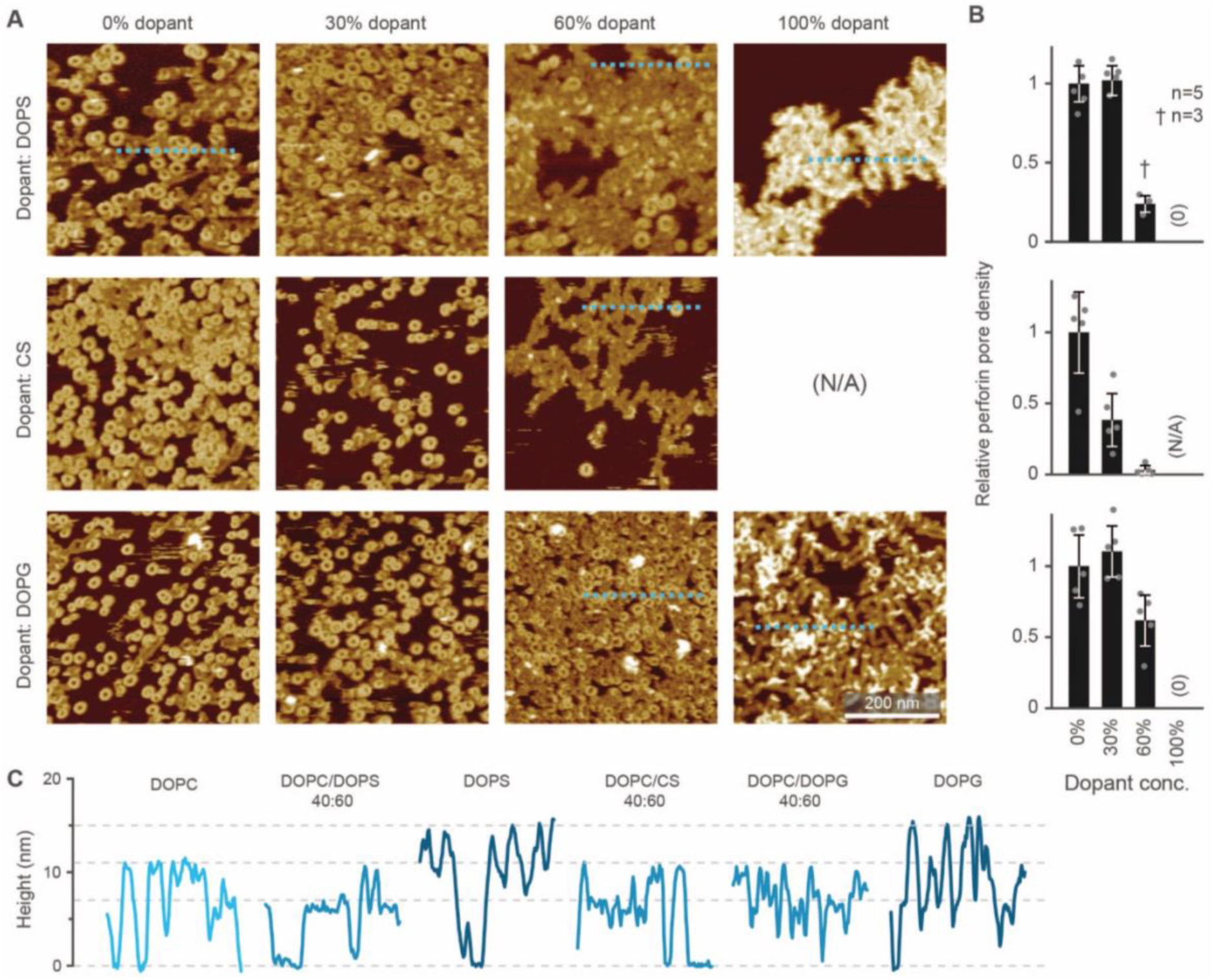
WT-PRF pore formation on different substrates containing DOPC and varying levels of either DOPS, DOPG, or cholesterol sulphate (CS). (A) Representative AFM images of perforin pores and aggregations on the different substrates. For pure CS, no bilayer could be formed. All data was recorded at room temperature. (B) Quantification of pore formation (mean ± SD) in the samples shown in (A), relative to the 0% dopant/ 100% DOPC reference. (C) Height profiles extracted along the dashed lines in A; the ‘DOPC’ reference profile was extracted from the first tile in A. The different profiles show the membrane level adjusted to 0 nm and the height of perforin pores (ca. 11 nm) and aggregates (ca. 7 nm at 60% dopant levels, up to 15 at 100% dopant levels), as highlighted by horizontal dashed lines. The colour tone of the profiles is darker compared to the ‘DOPC’ reference, relative to the negative charge present in the membrane substrates. Panels A and B are reproduced from Ref. ^14^, permission pending.

On the membranes with higher negative charge, the decrease in WT-PRF pore formation was accompanied by an increase in the presence of plaques of protein aggregates. Perhaps the most striking feature of these plaques is their height. When PC is present with at least 30 mol%, perforin aggregations appear at ca. 7 nm in height. However, on pure DOPS and DOPG membranes, the aggregations appear as plaques with a height of (up to) 15 nm above the membrane surface (Figure 6 C). This height is to be compared with the ca. 11 nm height of a membrane-bound perforin prior to and after membrane insertion ^13,14^.

Intriguingly, the observed 15 nm height above the membrane agrees with the full height of perforin pore including the hairpins that span the membrane ^7^. This suggests that the protein has initiated the transition from its prepore to pore state, yet in unfurling these hairpins has failed to insert into the membrane. Given this possible interpretation, we sought to first further validate the height measurements of perforin plaques on pure PS membranes, by including the CDC perfringolysin O (PFO) as a height ruler in our experiments. Like other CDCs, PFO forms pores that protrude ca. 7 nm above the membrane ^45,46,58,59^. To this end, we prepared DOPS/cholesterol 70:30 mol% membranes, on which perforin behaved similarly as on pure DOPS membranes (Figure 7 A) but which – by the inclusion of cholesterol – allowed CDC binding and pore formation too (Figure 7 B). By first incubating these membranes with perforin and next with the CDC PFO, we observed PFO pores in addition to perforin plaques (Figure 7 C, D), with the perforin plaques being of approximately double the height above the membrane as the PFO pores, which were taken as a height reference of ca. 7 nm (Figure 7 E). This fully confirms the extraordinary height of perforin on DOPS-only bilayers.

**Figure 7.**
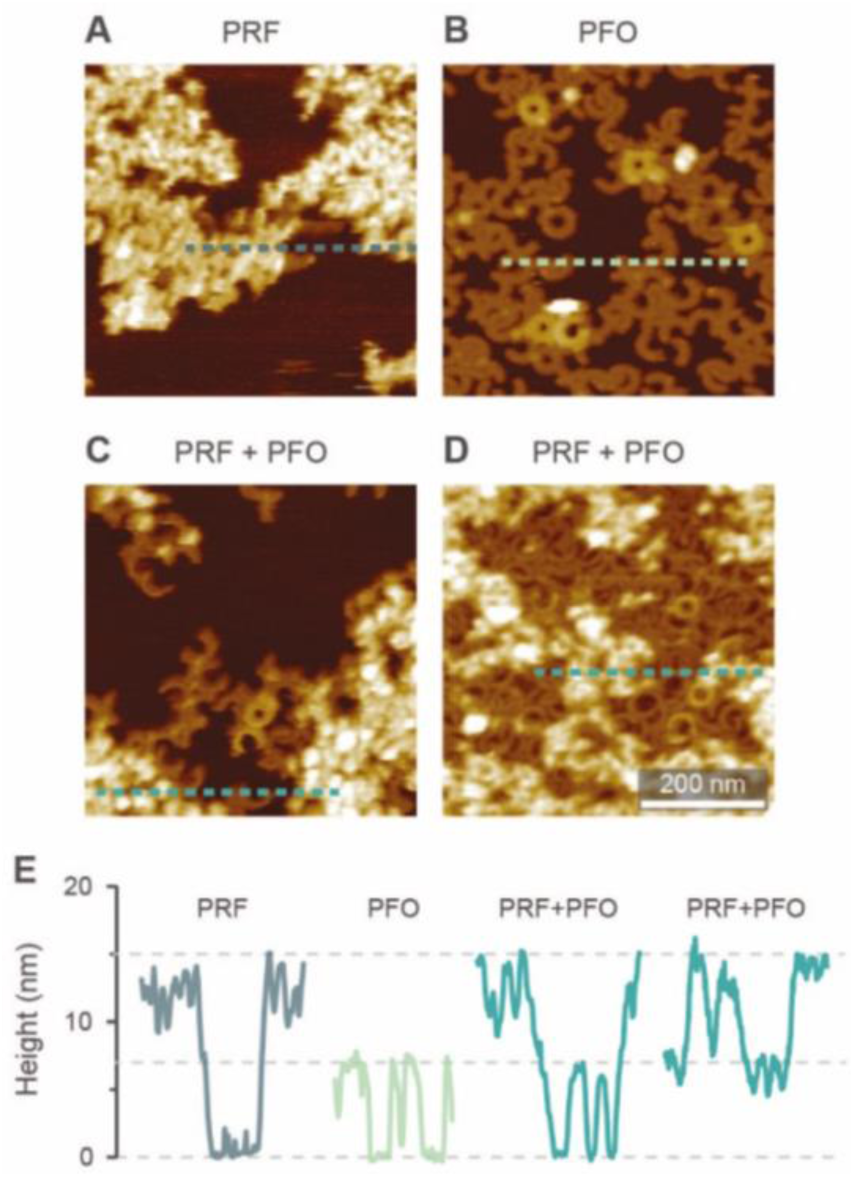
AFM images of WT-PRF and PFO on DOPS/cholesterol 70:30 mol% bilayers for height referencing. (A) WT-PRF forms protein plaques on the lipid bilayer. (B) Cholesterol dependent PFO pore formation, visible as arc- and ring-shaped assemblies. A small number of the assemblies appeared higher, probably due to incomplete membrane insertion: PFO collapses from ca. 10 nm to 7 nm height upon membrane insertion ^41^. (C, D) When the DOPS/cholesterol bilayers were first incubated with WT-PRF and next with PFO, perforin plaques were observed adjacent to PFO pores. We here show two samples incubated with different amounts of PFO, ca. 150 nM in C and ca. 450 nM in D. As a consequence, the membrane surface still is visible in C, while in D the PFO pores cover most of the remaining membrane. (E) Height profiles extracted along the dashed lines in A-D. Horizontal lines at 0, 7, and 15 nm highlight the membrane surface and the heights of PFO pores and perforin plaques, respectively. All AFM data were recorded at room temperature.

The preconditions and structural changes necessary to form such perforin plaques are unknown. In our earlier studies with the non-functional TMH1-PRF on pure DOPS bilayers, it emerged that in the membrane-binding and early assembly stage, *i.e*., before pore insertion, the behaviour of perforin is similar to that observed on DOPC bilayers^13,14^: TMH1-PRF on DOPS and DOPC (i) showed a similar distribution of subunits per assembly, (ii) was ca. 10 nm in height above the membrane, and (iii) was freely diffusing and could be removed from membrane surface by chelation of Ca^2+^ (demonstrated by the removal of oligomers after chelating calcium from the buffer). Only after adding DTT and thus unlocking TMH1-PRF, the short oligomers clustered together and increased their height to ca. 15 nm. Taken together, this supports the interpretation that the formation of plaques on PS is linked to the unfurling of the protein as it attempts – unsuccessfully – to insert into the membrane.

To further investigate how this behaviour depends on electrostatic interactions, we varied the concentration of divalent ions in solution, thus changing the screening of surface charges. Firstly, when 5 mM Ca^2+^ and an excess concentration of Mg^2+^ (25 mM) was present in the buffer, perforin (WT-PRF) would not bind to or form plaques on a DOPS membrane, even at higher perforin concentrations (Figure 8 A,B). Secondly, in the absence of Mg^2+^, the density of the plaques was dependent on the Ca^2+^ concentration in the buffer: the more Ca^2+^, the more condensed the formation of the plaques (Figure 8 C,D). The dependence of perforin membrane binding on the concentration of divalent cations further confirms that the observed behaviour on DOPS is mediated by electrostatic interactions. These two observations are different from what we observe on DOPC membranes, where a similar increase in Ca^2+^ concentration produced no effect on the formation of arc- and ring-shaped pores.

**Figure 8.**
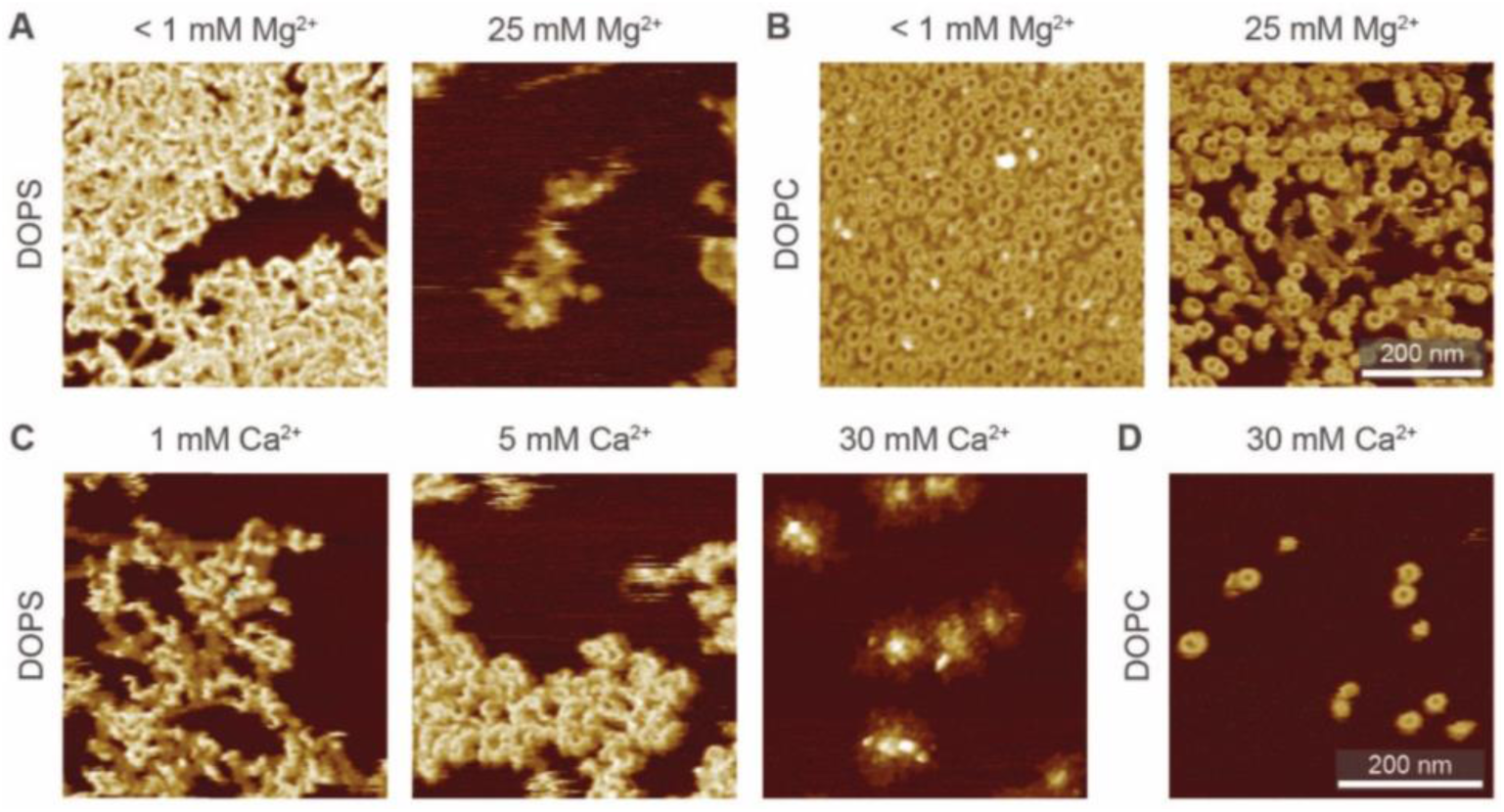
Interaction of WT-PRF with DOPS membranes at various levels of Ca^2+^ and Mg^2+^. (A) WT-PRF on DOPS membranes at low (< 1 mM) and high (25 mM) Mg^2+^ levels in the buffer. Only at low Mg^2+^ levels, perforin forms the ca. 15 nm high plaques, although some protein is binding at a high Mg^2+^ level. Here, we used 400 nM WT-PRF (instead of 150 nM for other experiments, see Methods) to test the effect at high perforin concentrations. (B) Analogous experiment to A on DOPC instead of DOPS membranes, as a control. At both low and high Mg^2+^ concentrations, arc- and ring-shaped perforin pores are visible. ^§§^ (C) Perforin plaques formed on DOPS membranes at Ca^2+^ concentrations of 1, 5, and 30 mM Ca^2+^ (and no Mg^2+^). For larger Ca^2+^ concentrations, the plaques appear more condensed and isolated. (D) For comparison, arc- and ring-shaped pores formed on DOPC at 30 mM Ca^2+^. All images were recorded at room temperature.

For functional perforin, the initial membrane binding occurs through its C2 domain. By mutating this domain, in D429A-PRF ^5^, we could also test whether the initial perforin binding depends on lipid composition: the mutation completely abrogated D429-PRF binding to DOPC bilayers, but on DOPS membranes the mutant still formed plaques with heights of mostly ca. 7 nm and up to 15 nm (Figure 9), roughly consistent with plaques formed by WT-PRF (Figure 6).

**Figure 9.**
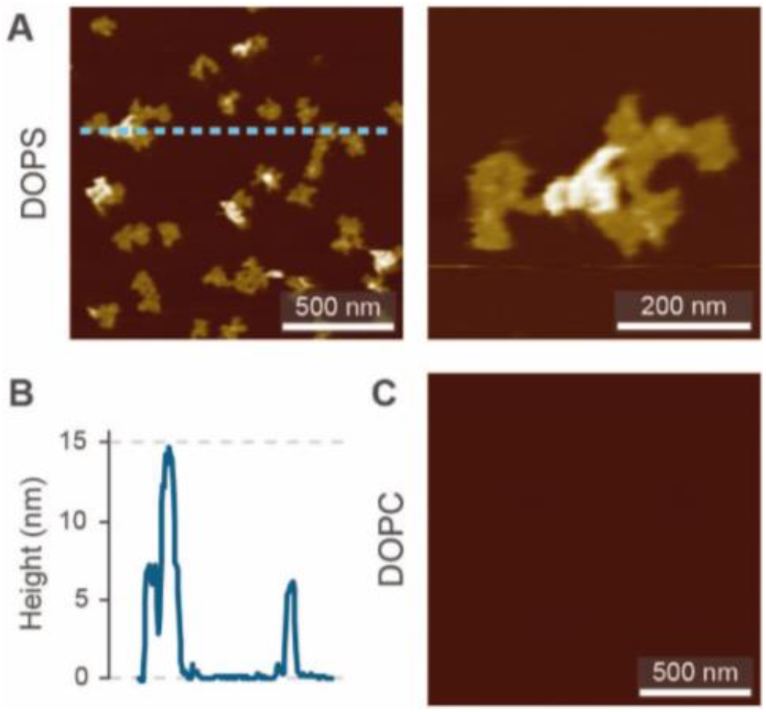
D429A-PRF, a perforin with mutated C2 domain, binds to DOPS but not DOPC membranes. (A) D429A-PRF forming plaques on a DOPS membrane. (B) Height profile extracted along the dashed line in A. (C) A DOPC membrane incubated with the same amount of D429A-PRF as in A does not show perforin binding. AFM data was recorded at room temperature.

Taken together, the experimental data indicate that negatively charged membranes disrupt perforin function due to electrostatic interactions at the stage of membrane insertion. Our data using D429A-PRF mutant indicate that such interactions take place at the interface between the membrane surface and parts of perforin that are not responsible for its functional Ca^2+^-dependent membrane binding through the C2 domain. One may therefore speculate that slight changes in the binding geometry of perforin eventually lead to protein misfolding and the formation of plaques on DOPS membranes. Another possibility is that only a fraction of perforin is required to bind DOPS in a different fashion, possibly independent of the C2 domain, and that this fraction disables otherwise correctly bound perforin when as it attempts to insert the membrane. Of note, a similar disruptive effect was observed when functional WT-PRF was co-incubated with excess non-functional TMH1-PRF ^13^, although in that case, pore forming functionality could be fully restored by subsequent addition of DTT (unlocking the disulphide lock in TMH1-PRF).

### Effect of membrane tension on perforin pore formation

Besides lipid order and charge^14^, another physical membrane property that may modulate perforin pore formation is membrane tension, which has been suggested to enhance perforin function in the immune synapse^60^. To some extent, such effects can be tested in supported lipid bilayers by the inclusion of curvature-inducing lipids. For example, phosphatidylethanolamine (PE) is a zwitterionic lipid with no net charge and a relatively small headgroup compared with the width of its hydrophobic tail. This causes PE to favour curved membrane arrangements, consistent with its prevalence in the inner leaflet of the eukaryotic plasma membrane and implying interfacial tension when forced to arrange in planar membranes; indeed, bilayers containing only (unsaturated) PE lipids do not form under physiological conditions ^61^. PE can be synthesized from PS by decarboxylation and co-locates with PS in the inner plasma membrane leaflet^62,63^; their externalization is regulated by the same transporters^64^.

To test how the addition of PE affects pore formation by perforin, we doped a DOPC bilayer with up to 60 mol% DOPE, and exposed the resulting membranes to WT-PRF. As shown in Figure 10 A,B, the addition of DOPE had no significant influence on the pore forming capacity of perforin *per se*. We also performed an alternative experiment in which we tested whether addition of 80 mol% DOPE would restore pore forming capacity in a DOPS host bilayer. A qualitative assessment of the AFM images indeed indicated that the inclusion of PE caused some restoration of pore formation on DOPS membranes (Figure 10 C). This preliminary result could be explained by presuming that perforin directly binds to PE lipid headgroups. However, it is also possible that PE, with its small headgroup, provides no direct perforin binding site and, instead, increases membrane tension and/or acts as a spacer between DOPS molecules. As a result, perforin might access and bind the PS headgroup differently, thus (partially) restoring its functionality. Future experiments will need to determine if perforin can bind PE directly.

**Figure 10.**
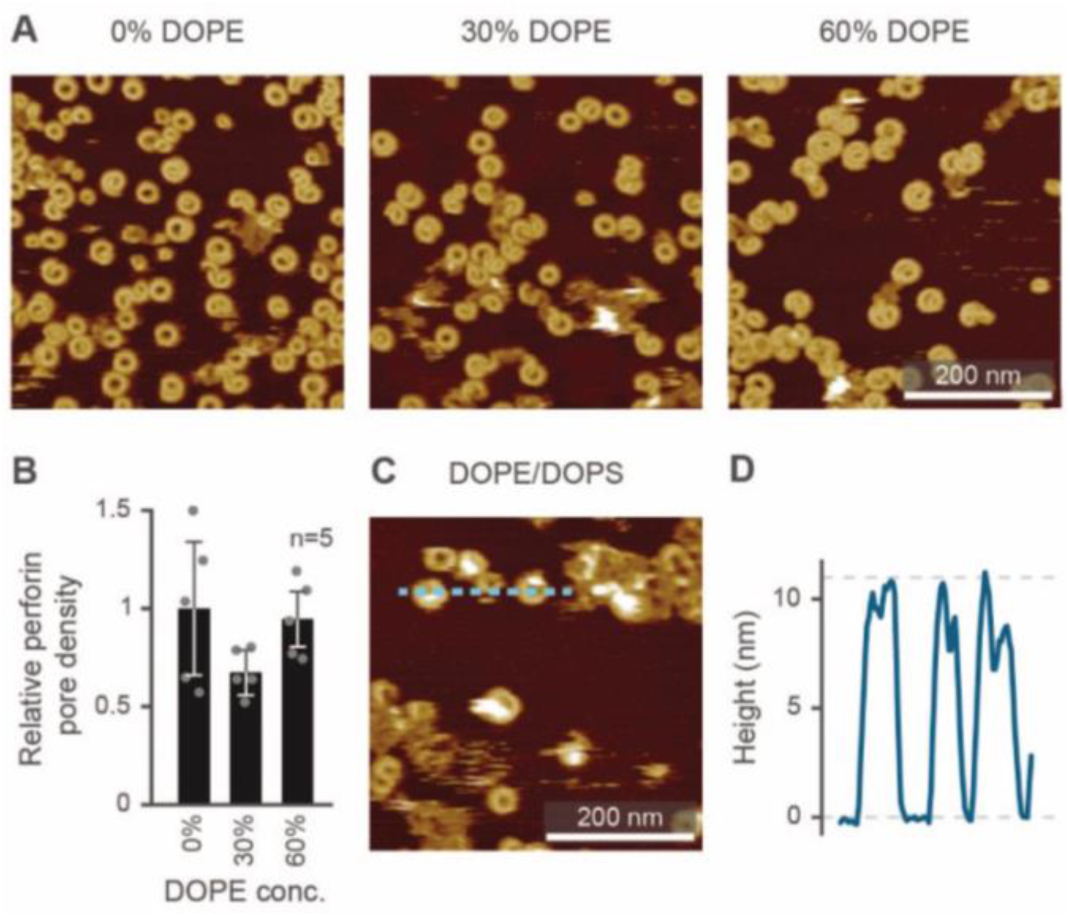
Effect of DOPE on WT-PRF pore formation. (A) AFM images of perforin pore formation on membranes containing DOPC and increasing amounts of DOPE. (B) Quantification of the pore formation normalized to the 0% DOPE/ 100% DOPC reference. Error bars represent standard deviations. (C) AFM image of perforin on a DOPS/DOPE 80:20 membrane, showing at least partial restoration of pore formation. AFM images were recorded at room temperature.

## Conclusions

As discussed in this paper, physical properties of membranes play essential roles in determining their sensitivity to perforin pore formation. This applies to the lipid order and packing, which reduce perforin binding to the membrane ^11,14^; to lipid charge, which causes perforin to be trapped in dysfunctional aggregates ^14^; and possibly to membrane tension, which has been reported to enhance perforin function in the immune synapse ^60^.

Compared with previous results, we have here (i) demonstrated the power of AFM and model membranes in investigating lipid specificity of pore forming proteins and of perforin in particular (ii) used AFM to demonstrate how membrane order in gel-phase lipids completely prevents perforin binding, as previously observed for liquid-ordered domains ^14^; (iii) demonstrated that this lipid specificity for liquid-disordered membranes is robust against variations in divalent ion concentration (Mg^2+^, and Ca^2+^ above the threshold needed to facilitate perforin binding to the membrane); (iv) validated the extraordinary height (ca. 15 nm above the membrane) of dysfunctional perforin aggregates observed on negatively charged membranes; (v) confirmed the electrostatic nature of how such membranes disable perforin; and (vi) showed that perforin pore formation is relatively insensitive to interfacial membrane tension, although it may play a role in rescuing perforin functionality on PS-rich membranes.

To assess the physiological relevance of these findings, they need to be compared with cell-based assays, e.g., possible correlations of perforin lysis with lipid order in target cell membranes ^12^, which confirm that reduction of lipid order sensitizes CTL membranes to perforin and that non-lytic perforin colocalizes with externalized (non-apoptotic) PS on CTLs ^14^.

Finally, it is noted that related pore forming proteins have been reported to show specificity for particular lipids, e.g., the membrane attack complex ^30^ and gasdermin ^31^ for negatively charged lipids, whereas bacterial CDCs, prefer cholesterol-rich and hence liquid ordered domains in phase-separated membranes ^46^. These observations indicate a wide range of biomedically relevant processes in which physical properties of membranes may be determinants for the function of pore forming proteins.

## Acknowledgements

We thank Richard Thorogate, Elena Taran, and Tian Zheng for technical support and access to AFM facilities; Peter Andrew and Rana Lonnen for providing PFO; Sandra Verschoor and Annette Ciccone for expression and purification of perforin and perforin mutants. This work has been funded by the NHMRC Project Grant (1128587), the NHMRC Fellowship (1059126), the BBSRC (BB/J005932/1, BB/J006254/1 and BB/N015487/1); the EPSRC (EP/M028100/1); the Sackler Foundation; and the Swiss National Science Foundation Grant p2skp3_187634.

## Notes and references

§ Some of the observed perforin-lipid binding was different from previously published results. Without investigating this further, we point out that we used a recombinant mouse perforin mutant (vs. native human perforin elsewhere) and detected fluorescence directly (vs. primary/secondary antibody detection elsewhere). Furthermore, we noted some perforin binding that exclusively occurred in the absence of Ca^2+^ and disappeared after prolonged incubation in the washing buffer. The reasons for this are unknown to us. §§ We note that in the images depicted here, overall perforin coverage at high Mg^2+^ concentrations appears lower (at 400 nM WT-PRF concentration). It is not clear if this difference is significant; to date, we do not have sufficient AFM data and repeats of this experiment to rigorously quantify the amount of perforin pores at high versus low Mg^2+^ concentrations.

## References

1 I. Voskoboinik, J. C. Whisstock and J. A. Trapani, Nat. Rev. Immunol., 2015, 15, 388–400.

2 P. Golstein and G. M. Griffiths, Nat. Rev. Immunol., 2018, 18, 527–535.

3 J. A. Lopez, M. R. Jenkins, J. A. Rudd-schmidt, A. J. Brennan, J. C. Danne, S. I. Mannering, J. A. Trapani and I. Voskoboinik, J. Immunol., 2013, 191, 2328–2334.

4 M. R. Jenkins, J. A. Rudd-Schmidt, J. A. Lopez, K. M. Ramsbottom, S. I. Mannering, D. M. Andrews, I. Voskoboinik and J. A. Trapani, J. Exp. Med., 2015, 212, 307–317.

5 I. Voskoboinik, M. C. Thia, J. Fletcher, A. Ciccone, K. Browne, M. J. Smyth and J. A. Trapani, J. Biol. Chem., 2005, 280, 8426–8434.

6 R. U. Moreno, J. Gil, C. Rodriguez-Sainz, E. Cela, V. LaFay, B. Oloizia, A. B. Herr, J. Sumegi, M. B. Jordan and K. A. Risma, Blood, 2009, 113, 338–346.

7 R. H. P. Law, N. Lukoyanova, I. Voskoboinik, T. T. Caradoc-Davies, K. Baran, M. A. Dunstone, M. E. D’Angelo, E. V Orlova, F. Coulibaly, S. Verschoor, K. a Browne, A. Ciccone, M. J. Kuiper, P. I. Bird, J. a Trapani, H. R. Saibil and J. C. Whisstock, Nature, 2010, 468, 447–51.

8 D. A. K. Traore, A. J. Brennan, R. H. P. Law, C. Dogovski, M. A. Perugini, N. Lukoyanova, E. W. W. Leung, R. S. Norton, J. A. Lopez, K. A. Browne, H. Yagita, G. J. Lloyd, A. Ciccone, S. Verschoor, J. A. Trapani, J. C. Whisstock and I. Voskoboinik, Biochem. J., 2013, 456, 323–335.

9 H. Yagi, P. J. Conroy, E. W. W. Leung, R. H. P. Law, J. A. Trapani, I. Voskoboinik, J. C. Whisstock and R. S. Norton, J. Biol. Chem., 2015, 290, 25213–26.

10 J. Tschopp, S. Schäfer, D. Masson, M. C. Peitsch and C. Heusser, Nature, 1989, 337, 272–274.

11 R. Antia, R. A. Schlegel and P. Williamson, Immunol. Lett., 1992, 32, 153–157.

12 D. M. Ojcius, S. Jiang, P. M. Persechini, J. Storch and J. D. Young, Mol. Immunol., 1990, 27, 839–845.

13 C. Leung, A. W. Hodel, A. J. Brennan, N. Lukoyanova, S. Tran, C. M. House, S. C. Kondos, J. C. Whisstock, M. A. Dunstone, J. A. Trapani, I. Voskoboinik, H. R. Saibil and B. W. Hoogenboom, Nat. Nanotechnol., 2017, 12, 467–473.

14 J. A. Rudd-Schmidt, A. W. Hodel, T. Noori, J. A. Lopez, H. J. Cho, S. Verschoor, A. Ciccone, J. A. Trapani, B. W. Hoogenboom and I. Voskoboinik, Nat. Commun., 2019, 10, 1–13.

15 N. Rojko and G. Anderluh, Acc. Chem. Res., 2015, 48, 3073–3079.

16 N. V. Dudkina, B. A. Spicer, C. F. Reboul, P. J. Conroy, N. Lukoyanova, H. Elmlund, R. H. P. Law, S. M. Ekkel, S. C. Kondos, R. J. A. Goode, G. Ramm, J. C. Whisstock, H. R. Saibil and M. A. Dunstone, Nat. Commun., 2016, 7, 10588.

17 N. Lukoyanova, B. W. Hoogenboom and H. R. Saibil, J. Cell Sci., 2016, 129, 2125–2133.

18 R. J. C. Gilbert, M. D. Serra, C. J. Froelich, M. I. Wallace and G. Anderluh, Trends Biochem. Sci., 2014, 39, 510–516.

19 C. F. Reboul, J. C. Whisstock and M. A. Dunstone, Biochim. Biophys. Acta - Biomembr., 2016, 1858, 475–486.

20 P. Drücker, I. Iacovache, S. Bachler, B. Zuber, E. B. Babiychuk, P. S. Dittrich and A. Draeger, Biomater. Sci., 2019, 7, 3693–3705.

21 B. B. Johnson, P. C. Moe, D. Wang, K. Rossi, B. L. Trigatti and A. P. Heuck, Biochemistry, 2012, 51, 3373–3382.

22 J. J. Flanagan, R. K. Tweten, A. E. Johnson and A. P, Biochemistry, 2009, 48, 3977–3987.

23 L. D. Nelson, A. E. Johnson and E. London, J. Biol. Chem., 2008, 283, 4632–4642.

24 J. P. Sáenz, D. Grosser, A. S. Bradley, T. J. Lagny, O. Lavrynenko, M. Broda and K. Simons, Proc. Natl. Acad. Sci. U. S. A., 2015, 112, 11971–11976.

25 E. J. Dufourc, J. Chem. Biol., 2008, 1, 63–77.

26 T. Tomita, K. Noguchi, H. Mimuro, F. Ukaji, K. Ito, N. Sugawara-Tomita and Y. Hashimoto, J. Biol. Chem., 2004, 279, 26975–26982.

27 K. Ota, A. Leonardi, M. Mikelj, M. Skočaj, T. Wohlschlager, M. Künzler, M. Aebi, M. Narat, I. Križaj, G. Anderluh, K. Sepčić and P. Maček, Biochimie, 2013, 95, 1855–64.

28 N. Lukoyanova, S. C. Kondos, I. Farabella, R. H. P. Law, C. F. Reboul, T. T. Caradoc-Davies, B. A. Spicer, O. Kleifeld, D. A. K. Traore, S. M. Ekkel, I. Voskoboinik, J. A. Trapani, T. Hatfaludi, K. Oliver, E. M. Hotze, R. K. Tweten, J. C. Whisstock, M. Topf, H. R. Saibil and M. A. Dunstone, PLOS Biol., 2015, 13, e1002049.

29 M. Novak, T. Krpan, A. Panevska, L. K. Shewell, C. J. Day, M. P. Jennings, G. Guella and K. Sepčić, Biochim. Biophys. Acta - Biomembr., 2020, 183307.

30 E. S. Parsons, G. J. Stanley, A. L. B. Pyne, A. W. Hodel, A. P. Nievergelt, A. Menny, A. R. Yon, A. Rowley, R. P. Richter, G. E. Fantner, D. Bubeck and B. W. Hoogenboom, Nat. Commun., 2019, 10, 1–10.

31 P. Broz, P. Pelegrín and F. Shao, Nat. Rev. Immunol., 2020, 20, 143–157.

32 R. N. Lewis, B. D. Sykes and R. N. McElhaney, Biochemistry, 1988, 27, 880–887.

33 K. Baran, M. Dunstone, J. Chia, A. Ciccone, K. A. Browne, C. J. P. Clarke, N. Lukoyanova, H. Saibil, J. C. Whisstock, I. Voskoboinik and J. A. Trapani, Immunity, 2009, 30, 684–695.

34 J. A. Lopez, A. J. Brennan, J. C. Whisstock, I. Voskoboinik and J. A. Trapani, Trends Immunol., 2012, 33, 406–412.

35 S. S. Metkar, M. Marchioretto, V. Antonini, L. Lunelli, B. Wang, R. J. C. Gilbert, G. Anderluh, R. Roth, M. Pooga, J. Pardo, J. E. Heuser, M. D. Serra and C. J. Froelich, Cell Death Differ., 2015, 22, 74–85.

36 R. N. Lewis, N. M. Nanette Mak and R. N. McElhaney, Biochemistry, 1987, 26, 6118–6126.

37 S. L. Veatch and S. L. Keller, Phys. Rev. Lett., 2005, 94, 3–6.

38 R. F. M. De Almeida, A. Fedorov and M. Prieto, Biophys. J., 2003, 85, 2406–2416.

39 K. Furuya and T. Mitsui, J. Phys. Soc. Japan, 1979, 46, 611–616.

40 R. A. Demel, F. Paltauf and H. Hauser, Biochemistry, 1987, 26, 8659–8665.

41 P. W. M. Van Dijck, BBA - Biomembr., 1979, 555, 89–101.

42 E. B. Smaal, K. Nicolay, J. G. Mandersloot, J. de Gier and B. de Kruijff, BBA - Biomembr., 1987, 897, 453–466.

43 N. Yilmaz and T. Kobayashi, Biochim. Biophys. Acta - Biomembr., 2016, 1858, 500–511.

44 A. W. Hodel, C. Leung, N. V. Dudkina, H. R. Saibil and B. W. Hoogenboom, Curr. Opin. Struct. Biol., 2016, 39, 8–15.

45 D. M. Czajkowsky, E. M. Hotze, Z. Shao and R. K. Tweten, EMBO J., 2004, 23, 3206–3215.

46 C. Leung, N. V Dudkina, N. Lukoyanova, A. W. Hodel, I. Farabella, A. P. Pandurangan, N. Jahan, M. Pires Damaso, D. Osmanović, C. F. Reboul, M. A. Dunstone, P. W. Andrew, R. Lonnen, M. Topf, H. R. Saibil and B. W. Hoogenboom, Elife, 2014, 3, e04247.

47 S. D. Connell and D. A. Smith, Mol. Membr. Biol., 2006, 23, 17–28.

48 V. R. Sutton, N. J. Waterhouse, K. Baran, K. Browne, I. Voskoboinik and J. A. Trapani, Methods, 2008, 44, 241–249.

49 M. J. Hope, M. B. Bally, G. Webb and P. R. Cullis, Biophys. Biopphysica Acta, 1985, 812, 55–65.

50 C. P. S. Tilcock, Chem. Phys. Lipids, 1986, 40, 109–125.

51 J. R. Kremer, D. N. Mastronarde and J. R. McIntosh, J. Struct. Biol., 1996, 116, 71–76.

52 T. Hothorn, F. Bretz and P. Westfall, Biometrical J., 2008, 50, 346–363.

53 A. J. Farrand, S. LaChapelle, E. M. Hotze, A. E. Johnson and R. K. Tweten, Proc. Natl. Acad. Sci., 2010, 107, 4341–4346.

54 X. Liu, Z. Zhang, J. Ruan, Y. Pan, V. G. Magupalli, H. Wu and J. Lieberman, Nature, 2016, 535, 153–158.

55 S. S. Pang, C. Bayly-Jones, M. Radjainia, B. A. Spicer, R. H. P. Law, A. W. Hodel, E. S. Parsons, S. M. Ekkel, P. J. Conroy, G. Ramm, H. Venugopal, P. I. Bird, B. W. Hoogenboom, I. Voskoboinik, Y. Gambin, E. Sierecki, M. A. Dunstone and J. C. Whisstock, Nat. Commun., 2019, 10, 1–9.

56 G. Pabst, A. Hodzic, J. Štrancar, S. Danner, M. Rappolt and P. Laggner, Biophys. J., 2007, 93, 2688–2696.

57 R. J. Elin, Disease-a-Month, 1988, 34, 166–218.

58 S. J. Tilley, E. V Orlova, R. J. C. Gilbert, P. W. Andrew and H. R. Saibil, Cell, 2005, 121, 247–256.

59 K. van Pee, E. Mulvihill, D. J. Müller and Ö. Yildiz, Nano Lett., 2016, 16, 7915–7924.

60 R. Basu, B. M. Whitlock, J. Husson, A. L. Floc’h, W. Jin, A. Oyler-Yaniv, F. Dotiwala, G. Giannone, C. Hivroz, N. Biais, J. Lieberman, L. C. Kam and M. Huse, Cell, 2016, 165, 100–110.

61 V. A. Frolov, A. V. Shnyrova and J. Zimmerberg, Cold Spring Harb. Perspect. Biol., 2011, 3, a004747.

62 G. van Meer, D. R. Voelker and G. W. Feigenson, Nat. Rev. Mol. Cell Biol., 2008, 9, 112–124.

63 P. A. Leventis and S. Grinstein, Annu. Rev. Biophys., 2010, 39, 407–427.

64 J. H. Stafford and P. E. Thorpe, Neoplasia, 2011, 13, 299–308.

